# Disease Resistance correlates with Core Microbiome Diversity in Cotton

**DOI:** 10.1101/2023.08.25.554784

**Authors:** Rhea Aqueel, Ayesha Badar, Nazish Roy, Umer Zeeshan Ijaz, Kauser Abdulla Malik

**Author notes:** Joint corresponding authors (both authors jointly directed this work) (http://userweb.eng.gla.ac.uk/umer.ijaz).

## Abstract

Understanding the resident microbial communities and their above and below ground interactions with plants will provide necessary information for crop disease protection and stress management. In this study, we show how diversity of core microbiome varies with disease susceptibility of a crop. To test this hypothesis, we have focused on identifying the core microbial species of Cotton Leaf Curl Disease (CLCuD) susceptible *Gossypium hirsutum* and CLCuD resistant *Gossypium arboreum* under viral infestation. Derivation of core membership is challenging as it depends on an occupancy threshold of microbial species in a sampling pool, whilst accounting for different plant compartments. We have used an abundance-occupancy distribution approach where we dynamically assess the threshold for core membership, whilst marginalizing for occupancy in four compartments of the cotton plant, namely, leaf epiphyte, leaf endophyte, rhizosphere, and root endophyte. Additionally, we also fit a neutral model to the returned core species to split them into three groups, those that are neutral, those that are selected by the plant environment, and finally those that are dispersal limited. We have found strong inverse relationship between diversity of core microbiome and disease susceptibility. A deeper understanding of this association will aid in the development of biocontrol agents for improving plant immunity against biotrophic pathogens.

## Main

Microbes are minute but essential components of the environment which play crucial roles in the host plant’s response to disease. Insights into plant-microbe interactions have revealed a great deal about the underlying mechanisms that contribute to plant health and development. Apart from exploration of the entire microbiome, recent approaches have revealed interesting insights relating to the microbes that make up the core of plants and influence functional relationships with the host (Wang *et al*., 2023). Core microbial communities are the smallest subset of stable taxa identified in plant ecosystems, that can aid the plant in nutrient and water uptake, and promotes plant health by activating defense responses against biotic and abiotic stresses (Compant *et al*., 2019).

Past attempts at identifying core microbiome membership relied on a strict occupancy/prevalence threshold (Shetty *et al*., 2017) that varied between 50% to 95% depending on a study in consideration, which is always a source of debate. To circumvent this, Shade and Stopnisek (2019) have proposed a dynamic approach where microbial species are first ranked by their occupancy according to the considered study design by using both *site-specific occupancy*, and *replicate consistency* of microbial species. Starting with a seed core subset of top-ranked species, contribution of the core set to beta diversity is calculated, with species iteratively added until adding one more species offers diminishing returns on explanatory value for beta diversity. Thus, the occupancy threshold is learnt from the dataset. Furthermore, the approach is merged with Burns *et al*., (2016) approach, where neutral model is applied to species-occupancy distribution of observed microbial species. The 95% confidence intervals of neutral models are obtained for species plotted by mean log10 relative abundance and occupancy. The species that fall outside the 95% model are then inferred to be deterministically, rather than neutrally, selected. Overlapping the two approaches then provides a consistent methodology for prioritizing ecologically important core microbial species over space and time.

We have used the above core microbiome approach (Shade and Stopnisek, 2019) on an Amplicon Sequence Variants (ASVs) abundance table (n=59 samples x P=38,120 ASVs) from Aqueel *et al*., (2023). After pre-processing for low read samples and excluding contaminants, a total of 50 samples were retained with 34,144 ASVs. The samples were obtained for three varieties with varying susceptibility to Cotton Leaf Curl Disease (CLCuD): *Gossypium hirsutum* susceptible variety (PFV-2); *Gossypium hirsutum* partially tolerant variety (PFV-1); and *Gossypium arboreum* tolerant variety (FDH-228). Furthermore, the compartments for each variety under study were leaf epiphyte, leaf endophyte, rhizosphere, and root endophyte, and have therefore used the compartment-specific occupancy model. The resultant core microbiome is shown in Figures 1 and 2.

**Figure 1:**
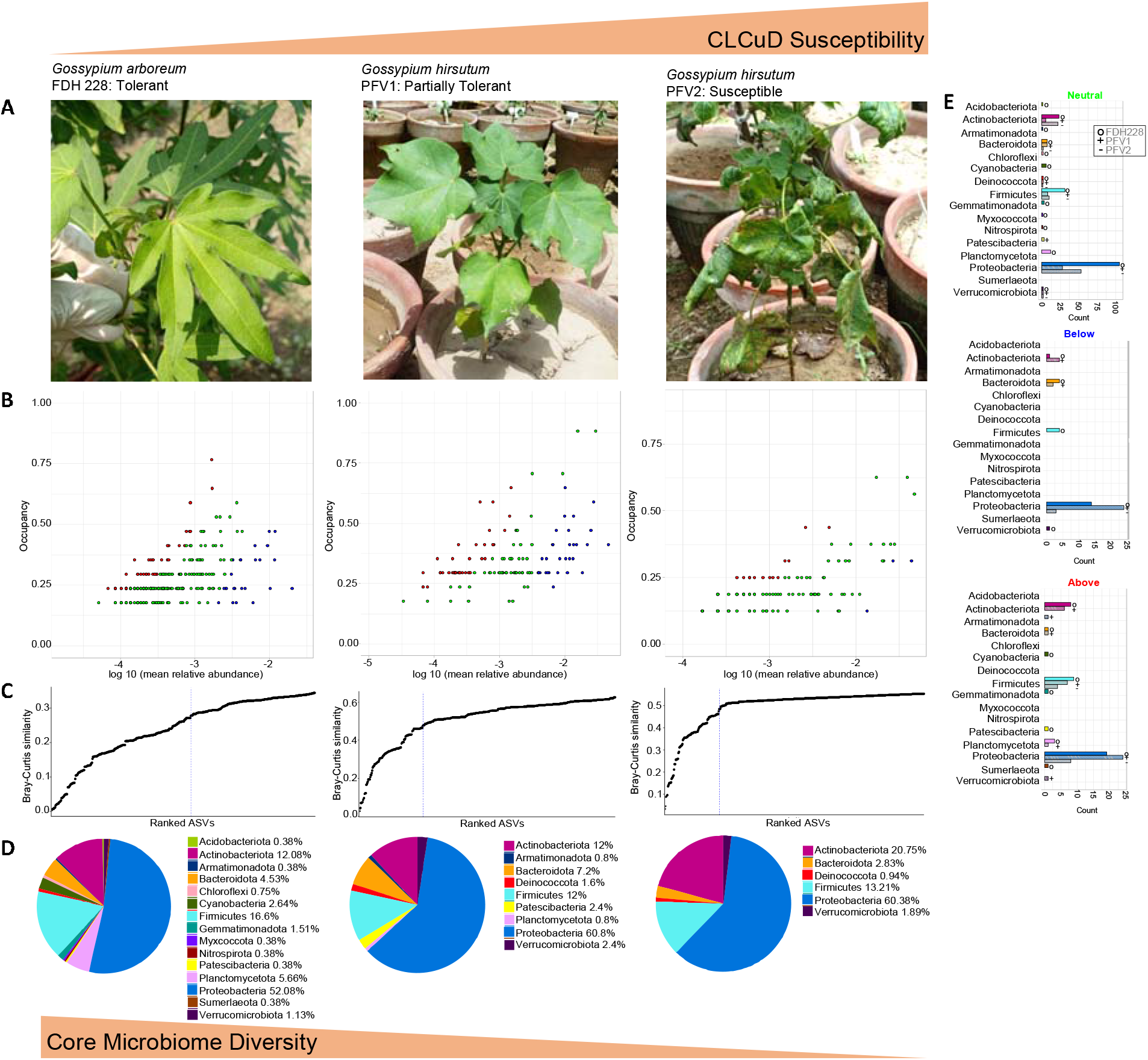
Core microbiome for three CLCuD varieties are shown in (A) and identified through species occupancy abundance diagrams in (B) incorporating a *Plant Compartment Specific Occupancy* model (compartments being *Leaf Epiphyte, Leaf Endophyte, Rhizosphere*, and *Root Endophyte*). Once we have obtained the ASV rankings depending on ASVs occupancy within these compartments as well as their replicate consistency, Bray-Curtis similarity is calculated for the whole dataset, and then also for only the top-ranked taxa. The contribution of the top-ranked taxa is divided by the total Bray-Curtis similarity to calculate a percent contribution of the prospective core set to beta diversity. The next-ranked taxon is added consecutively to find the point in the ranking at which adding one more taxon offers diminishing returns on explanatory value for beta diversity, and is shown in (C). The blue dotted line represents the “Last 2% decrease” criteria where ASVs are incorporated in the core subset until there is no more than 2% decrease in beta diversity. (D) represents the Phylum level assignment of the ASVs. Independently a neutral model is fitted with those ASVs that fall within the 95% confidence interval, shown in green in (B), and those that fall outside the 95% model confidence to be inferred as deterministically assembled, i.e., non-neutral ASVs. Points above the model are selected by the host environment, shown in red in (B), and points below the model are dispersal limited, shown in blue in (B). The count of neutral/non-neutral ASVs at Phylum level are shown with the bar plots in (E).

**Figure 2:**
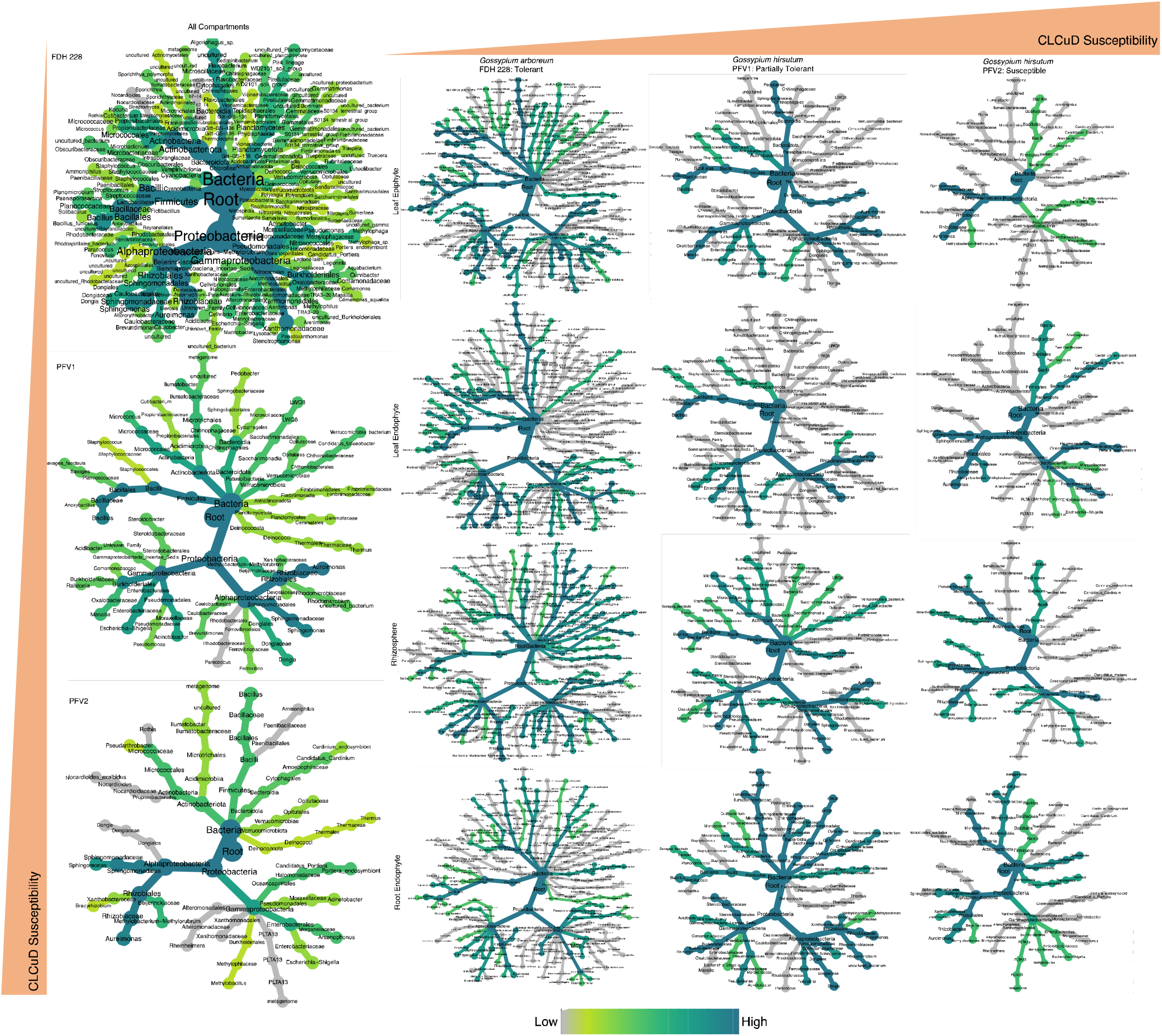
Taxonomic coverage of core microbiome in CLCuD susceptible and tolerant varieties, as calculated in Figure 1. Left (all compartments) Right (Separate coverage in different compartments: Leaf Epiphyte, Leaf Endophyte, Rhizosphere, and Root Endophyte).

The core microbiome has differentiating patterns in terms of disease resistance exhibiting higher diversity in the CLCuD resistant FDH-228 as compared to the partially susceptible PFV-1 and susceptible PFV-2. The distinctive phyla such as Sumerlaeota, Myxococcota, Gemmatimonadota, Patescibacteria, Deinococcota, Chloroflexi and Nitrospirota were identified in the core microbiome of FDH-228. Our findings support that CLCuD resistant variety associated microbiome is dynamic and is therefore selected by the cotton species itself due to its underlying functional relationships with the host plant.

Our targeted sequencing effort gives a precise insight into the core microbial communities of the susceptible, partially tolerant, and tolerant cotton varieties infected with the cotton leaf curl virus. On the phylum level, the different plant species under CLCuD attack had specific effects on the bacterial communities. The core taxa exhibited varying shifts between *Gossypium hirsutum* and *Gossypium arboreum*. Although, there were a few ASVs shared between the microbiomes, however, phyla such as Sumerlaeota and Gemmatimonadota which are known to thrive in harsh environments, were only identified in the CLCuD resistant FDH-228. Members of Gemmatimonadota are known to withstand harsh environments as saline soils and form a very small fraction of the bacterial community (Mujakić *et al*., 2022). The plant susceptibility to viruses is exhibited in the diversity of the core microbiome. Patescibacteria were linked with the partially tolerant PFV-1 and tolerant FDH-228. A study found Acidobacteriota in the core taxa of the interspecific interaction zone of peanut and sorghum rhizosphere (Shi *et al*., 2021). Acidobacteriota has only been identified in the core microbial taxa of the CLCuD resistant FDH-228. After fitting the neutral model, as compared to the susceptible varieties PFV-1 and PFV-2, we have found FDH-228 to be selecting more for Actinobacteriota, Cyanobacteria, Firmicutes, Gemmatimonadota, Patescibacteria, Planctomycetota, and Sumerlaeota. Actinobacteriota, Patescibacteria, and Planctomycetota, have been reported in the root endosphere of *Myrothamnus flabellifolia* which can withstand extreme drought conditions (Tebele *et al*., 2023).

CLCuD is one of the major biotic stresses in Pakistan causing severe economic losses up to more than US$2 billion per annum (Rafiq *et al*., 2021) in cotton crops, and with Pakistan being one of the top cotton producers, any strategy to suppress CLCuD will have significant impacts. The observed symptoms of CLCuD include vein thickening, leaf curling, leaf enation and dwarfing in the highly susceptible cotton genotypes eventually lead to mortality of the plant (Afzal *et al*., 2023; Vij *et al*., 2022; Pan *et al*., 2018; Sattar *et al*., 2013). There is very little (Aqueel *et al*., 2023) or no evidence to suggest that the use of plant-associated microbiome will lead to suppression of CLCuD. Whilst the diploid *Gossypium arboreum* harbors resistance genes against CLCuD, the widely cultivated *Gossypium hirsutum* remains susceptible. Treatment with the members of core microbial community inferred from the *Gossypium arboreum* may hold the key to safeguard *Gossypium hirsutum* which is cultivated >90% world wide (Li *et al*., 2015). Rhizospheric core microbial communities are previously known to offer promising plant disease resistance (Durán *et al*., 2018; Berendsen *et al*., 2012), albeit for *Arabidopsis*. Nonetheless, the construction of synthetic communities (SynComs) remains challenging, and depends on several factors including the spatial and temporal dynamics and also on the coordination of the microbes forming the community (Li *et al*., 2019; Johns *et al*., 2016). Through this research, and using a dynamic inferential approach, we anticipate that the obtained core microbiome of *Gossypium arboreum* (FDH-228: tolerant) and its diversity will aid in developing SynComs and may offer biocontrol potential by altering the plant-microbe ecology in the CLCuD infected cotton plant.

## Data availability

The datasets generated and analyzed during the current study are available from the corresponding author on reasonable request.

## Supplementary Information

The detailed dataset for Figure 1 and Figure 2 is provided as Data_Table_S1.xlsx

## Acknowledgments

We would like to thank all the field and laboratory teams of Kauser Abdullah Malik School of Life Sciences. We are grateful to Four Brothers Research Group, Pakistan for aiding us with the sampling of plants for this study.

## Author information

### Contribution

Rhea Aqueel (Conceptualization, Visualization, Data Curation, Investigation, Formal analysis, Writing - Original Draft)

Ayesha Badar (Data Curation, Visualization, Formal analysis, Investigation, Writing - Review & Editing)

Nazish Roy (Conceptualization, Methodology, Writing - Review & Editing, Supervision, Funding Acquisition)

Umer Zeeshan Ijaz (Methodology, Software, Formal analysis, Writing - Original Draft, Supervision, Funding Acquisition)

Kauser Abdulla Malik (Conceptualization, Methodology, Writing – review & editing, Project Administration, Supervision, Funding Acquisition)

